# Integrative Imaging of Lung Micro Structure: Amplifying Classical Histology by Paraffin Block μCT and same-slide Scanning Electron Microscopy

**DOI:** 10.1101/2024.06.29.601332

**Authors:** Johanna Reiser, Jonas Albers, Angelika Svetlove, Mara Mertiny, Felix K.F. Kommoss, Constantin Schwab, Anna Schneemann, Giuliana Tromba, Irene Wacker, Ronald E. Curticean, Rasmus R. Schroeder, Hans-Ulrich Kauczor, Mark O. Wielpütz, Christian Dullin, Willi L. Wagner

**Affiliations:** University Hospital Heidelberg, Dept. Diagnostic and Interventional Radiology, Im Neuenheimer Feld 420, 69120 Heidelberg, Germany; Biological X-ray Imaging, European Molecular Biology Laboratory, Hamburg Unit c/o DESY, Hamburg, Germany; Institute of Pathology, University Hospital of Heidelberg, Heidelberg, Germany; Carl Zeiss Microscopy GmbH, Carl-Zeiss-Strasse 22, 73447 Oberkochen, Germany; Italian synchrotron “Elettra”, Strada Statale 14 - km 163,5 in AREA Science Park, 34149 Basovizza, Trieste, Italy; 3DMM2O, Cluster of Excellence (EXC-2082/1–390761711) and Cryo Electron Microscopy, BioQuant, Heidelberg University and University Hospital, Heidelberg, Germany; University Medicine Goettingen, Dept. Diagnostic and Interventional Radiology, Robert-Koch-Str. 40, 37075, Goettingen, Germany; MPI for Multidisciplinary Sciences - City Campus, Translational Molecular Imaging, Herman Rein Str. 3, 37075 Goettingen, Germany

**Keywords:** propagation based imaging, phase contrast, virtual histology, FFPE

## Abstract

Classical histopathology of formalin fixed and paraffin embedded (FFPE) tissue using light microscopy (LM) remains the undisputed gold standard in biomedical microstructural lung tissue analysis. To extend this method, we developed an integrative imaging and processing pipeline which adds 3D context and screening capabilities by micro-CT (μCT) imaging of the entire paraffin block and adds ultrastructural information by correlative same-slide scanning electron microscopy (SEM). The different modalities are integrated by elastic registration to provide hybrid image datasets.

Without compromising standard light microscopic readout, we overcome the limitations of conventional histology by combining and integrating several imaging modalities. The biochemical information contained in histological and immunohistological tissue staining is embedded into the 3D tissue configuration and is amplified by adding ultrastructural visualization of features of interest. By combining μCT and conventional histological processing, specimens can be screened, and specifically preselected areas of interest can be targeted in the subsequent sectioning process.

While most of the μCT data shown in the manuscript was acquired at a Synchrotron, we further demonstrate that our workflow can also by applied using X-ray microscopy.

## Introduction

Understanding the intricate structures and composition of soft tissues is a pivotal pursuit in various fields, from medicine to biology. Histology, the primary method employed for microstructural tissue analysis, offers unparalleled versatility and specificity, aided by various staining protocols and immunohistochemistry. However, histological analysis has inherent limitations, including its intrinsic restriction to 2D assessments, the limited spatial resolution of light microscopy and challenges in identifying and targeting specific sites of interest during tissue processing. The latter often necessitates labor-intensive serial sectioning, diminishing the diagnostic value, particularly in diseases with small lesions and high tissue heterogeneity. The lung is a prime example of an organ where 3D analysis adds significant value in the understanding of the complex structural integrity of pulmonary tissue. The demands on the architecture of the lung are enormous. The largest organ in the human body is subject to permanent life-long volume changes while its internal structure must allow respiratory air to be evenly distributed and to get as close as possible to capillary blood vessels to allow for efficient gas-exchange. On the micro-scale, this is achieved by the creation of self-stabilizing tensegrity structures, utilizing an absolute minimum of tissue substance. At the center of these structures are extracellular helical line elements that form alveolar entrance rings and balance alveolar surface tension to prevent alveolar collapse (Weibel, 2009). Pulmonary injuries or diseases often involve impairment of the delicate lung structure with damage patterns at different length scales, ranging from the macroscopy of the whole organ down to the sub-cellular level (Leonard-Duke et al., 2020). Those structural alterations are often scattered and heterogeneously distributed within the lung (Rutting et al., 2021). Thus, a major challenge in microstructural lung tissue analysis is to identify sites of interest prior to executing subsequent high-resolution analysis by microscopic techniques (D’Amico et al., 2024).

Different and sometimes even adverse preparation steps are usually employed for different imaging modalities to investigate a variety of length of scales, such as micro computed tomography (μCT), light microscopy (LM) and electron microscopy (EM). As a result, the hierarchical context is often lost in multi-scale and multi-modality imaging approaches. In this report, we present a novel workflow for the integrative analysis of standard formalin-fixed and paraffin embedded (FFPE) lung tissue by combining high-resolution μCT with guided microtome sectioning, subsequent classical histology and same-slide correlative scanning electron microscopy (SEM). In order to demonstrate that our approach is not limited to the use of Synchrotron based μCT system, we show that X-ray microscopy can also be used. μCT displays tissue structures in their 3D context, allowing for volumetric analysis and serving as a screening tool for targeted sub-biopsies and guided microtome sectioning (Albers et al., 2018). Subsequent standard histology allows for the classification of cell types via specific staining, and tissue components can be integrated into their 3D context by elastic registration (Albers et al., 2021). Correlative same-slide SEM then provides insight into the ultrastructure of features of interest (Hyams et al., 2020), completing a multi-scale approach that can be applied to any unstained FFPE tissue, the persistent gold standard for biomedical tissue preparation and archiving. The integration of further additional modalities for downstream analysis of tissue sections and its integration into the 3D context are to be explored.

## Results

### Paraffin block μCT enhances comprehensive understanding of pulmonary soft tissues

The novel workflow that was applied uses unstained FFPE porcine lung tissue. Due to the elevated soft-tissue contrast of phase contrast CT acquired with synchrotron radiation (SRμCT) no additional staining was needed. This implies that the workflow can be applied to any unstained FFPE samples of soft tissues. It allows not only guided histological sectioning and targeted histopathological analysis but also for a comprehensive understanding of structures of interest in their three-dimensional context combined with information on their biochemical origin and their ultrastructure at a sub-micron resolution. Entire paraffin blocks of approximately 3×2×1 cm^3^ were scanned using the white beam setup of the SYRMEP beamline at the Italian Synchrotron light source “Elettra” (Dullin et al., 2021). Multiple scans with an isotropic voxel size of 4.5 μm were used to cover a tissue volume of 8×8×10 mm^3^, which allowed an overview of the tissue architecture at hand (Fig. 1A left). While information about the general tissue architecture could be obtained at this resolution, analysis of peripheral fine structures that become relevant at the micron-scale, like alveolar entrance rings, was not feasible (Fig. 1B). To obtain a scan at higher resolution with as little artifacts as possible, punch biopsies with a diameter of 4 mm were taken at preselected regions of interest and scanned again with 2 μm voxel size to form the base for subsequent 3D tissue analysis and identification of relevant structures (Fig. 1A right). Examination of these scans revealed a more detailed visualization of the general structure of pulmonary tissue (Fig. 1C). A higher level in soft tissue contrast was achieved, allowing the imaging of the complex 3D structure of the lung’s axial connective tissue system (Fig. 1D) which shapes the alveolar duct by forming alveolar entrance rings originating from bronchoalveolar duct junctions. To complete the workflow, microtome sectioning at identified sites of interest was performed. The acquired histological specimens were then examined by light microscopy and same-slide SEM.

**Figure 1.**
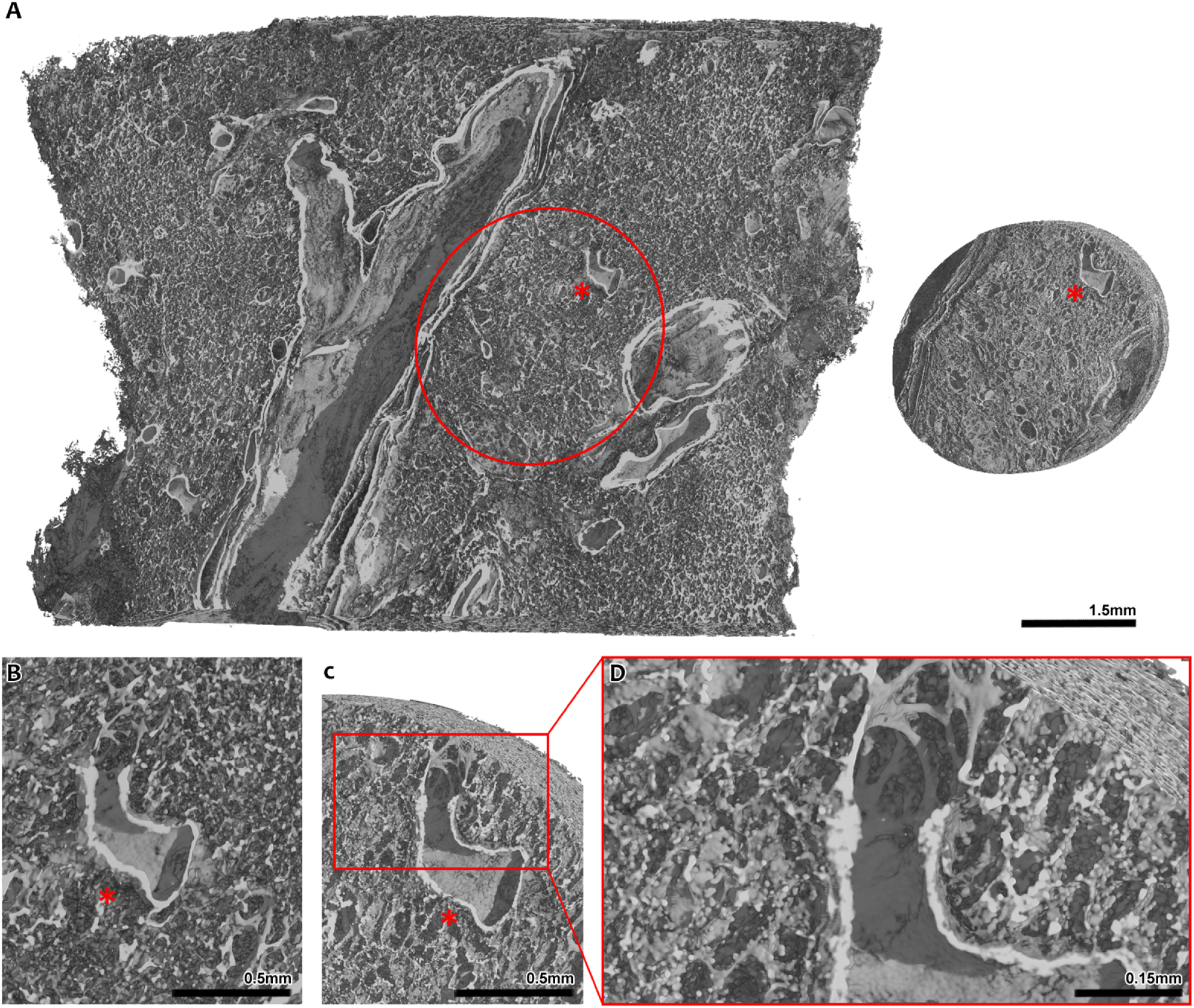
Comparison of synchrotron phase contrast SRμCT scans at different resolutions and fields of view. A) The volume shown on the left is a FFPE lung tissue block scanned at a voxel size of 4.5 μm. A cylindrical punch biopsy with a diameter of 4 mm was taken at the site marked in the volume on the left. On the right, said tissue cylinder is shown, scanned with a resolution of 2 μm. * marks a bronchoalveolar duct junction that, due to its unique shape, aids in correlating both scans and relocating one volume within the other. B) A zoomed in view of the bronchoalveolar duct junction (*) in the 4 μm resolution scan is shown. While its continuation into the alveolar space is visible, single structures like continuous interalveolar septa cannot be imaged at this resolution. C) A zoomed in view of the same bronchoalveolar duct junction (*) within the 2 μm biopsy cylinder scan. In direct comparison to C), details are depicted more clearly and small structural tissue details, like alveolar entrance rings, are captured due to the higher resolution of the scan. D) Detailed view of the alveolar entrance rings imaged in the 2 μm scan. Viewing the 3D rendering of the scan suggests that there is a connected network of these structures continuing inside the volume.

### μCT-guided screening and targeted microtome sectioning yields precise specific histological samples

To facilitate efficient μCT-guided histological sectioning, a correlation between modalities had to be established. Before microtome sectioning, the SRμCT scans were evaluated for potential sectioning planes of interest, containing structures relevant for histological correlation and analysis and prime quality with minimum tissue distortion and without air inclusions or fissures inside the paraffin. To determine the amount of tissue that had to be sectioned before reaching the desired planes of interest, 2-3 sections were first taken from the very top of the paraffin block and identified within the CT scan. This allowed calculation of the distance between the first sections close to the surface and the targeted sectioning plane of interest. According to these calculations, further sectioning was performed in several test runs, striking the specific plane of interest with an accuracy of ±14.45% relative to the calculated required depth. This deviation is attributed to the manual cutting process during which the FFPE sample is mounted and removed from the microtome several times for intermittent cooling. To account for the variation in the achieved cutting depth, tissue sections were not taken solely at a single targeted cutting plane but rather from a target zone. The extent of this target zone was relative to the cutting depth required to reach the desired sampling plane, spanning an additional 15% of this required depth in each direction of the cylinder. Using this refined method, acquired histological sections were then correlated back to the SRμCT scan to demonstrate the efficacy of the μCT-based targeted sectioning approach.

### Integrated Imaging can be achieved in a laboratory setting using X-ray microscopy devices

Synchrotron phase contrast SRμCT holds great advantages for this study as the high brilliance of the X-ray beam enables rapid scanning and the soft tissue contrast is enhanced by phase contrast effects. However, synchrotron radiation-based imaging is limited in accessibility. Thus, more accessible X-ray microscopy systems were explored and proved to provide sufficient spatial resolution and contrast to noise level to facilitate integrative imaging. In Fig. 2, the imaging results for a Zeiss Versa 620 (A) and the SYRMEP white beam setup (B) applied to the same porcine lung tissue specimen are compared. Zeiss Versa images demonstrate an even higher level of detail due to the fact that the voxel size at the detail scan was 1 μm, 2 times smaller than the voxel size of 2 μm within the SYRMEP data set. While the data set appears noisier, the lung tissue can clearly be separated from the paraffin embedding medium. Note the smooth appearance of the SYRMEP data sets is a result of the low pass filtering effect of the applied phase retrieval algorithm (Paganin et al., 2002). Thus, the data confirms that the attenuation contrast based X-ray microscope (Zeiss Versa 620) can be used to study unstained FFPE lung tissue specimens on a micron level. However, the acquisition time used for the data set shown in Fig. 6 (A) was 9.3 hrs. While this can certainly be optimized further, the high X-ray fluxes at Synchrotron sources typically allow very short acquisition times. In this study, the acquisition time per SRμCT dataset was only 180 s.

**Figure 2.**
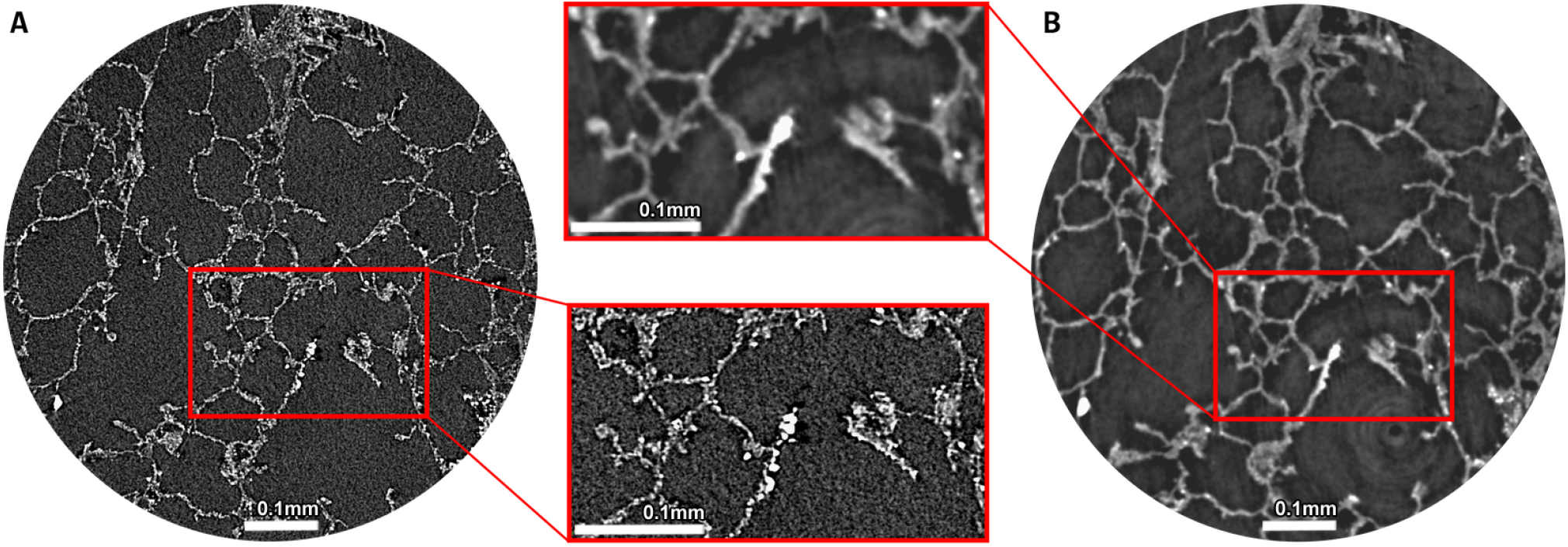
Comparison of attenuation based X-ray microscopy (Zeiss Versa 620 A) and phase contrast SRμCT (SYRMEP beamline B). As indicated by the detail views zooming in on a high-contrast septal tip, both classical X-ray microscopy and SRμCT can successfully be performed with unstained lung tissue embedded in paraffin. There is, however, a notable difference in scanning times with the settings applied in this study (9.3 hrs for Versa 620, 180s for SRμCT). Note: scans have been performed with different voxel sizes (1 μm Versa, 2 μm SYRMEP).

### Elastic image registration of LM and μCT allows for segmentation and integration of complex structural hallmarks of the peripheral lung

Despite the high-resolution 3D imaging capability of μCT, the unspecific contrast hinders precise classification of tissue components. Histology on the other hand allows for specific identification of various tissue types but can hardly provide 3D structural information. To overcome these limitations, we employed elastic image registration to co-register subsequently obtained and AB-PAS-stained lung tissue slides to the 3D context of the original uncut paraffin block. Fig. 3A) shows a defined virtual section through the SRμCT data set of an exemplary 4 mm diameter punch biopsy of an embedded porcine lung sample. Fig. 3B) shows the AB-PAS-stained histology slices using the SRμCT-derived cutting depth for targeted sectioning. Clearly, a strong resemblance between the virtual section in Fig. 3A) and the histological slide in Fig. 3B) can be appreciated. Since microtome sectioning causes non-uniform local deformations, elastic image registration was employed to register the histological slide to the corresponding virtual section within the SRμCT scan. Fig. 3C) demonstrates that the SRμCT data set can be used to access the local 3D environment of the structures seen in the histological slide. However, identification and reliable classification of certain tissue types is difficult because there is little data available for reference. Coregistration of the 3D SRμCT dataset and the digitalized micrograph generates a hybrid dataset (Fig. 3D). Complex structures, such as the broncho-alveolar-duct-junction (BADJ), can now be identified using the specific information from the digitalized light micrograph and its intricate micro-architecture can be assessed in 3D, portraying the convoluted course of the axial connective tissue system originating from the BADJ, giving rise to the backbone of the alveolar duct by forming the alveolar entrance rings (Fig. 3E). Tiny cross-sections of the cable-like structure can be appreciated in light microscopy only (black arrows in LM insert Fig. 3E). In the SRμCT data, a carina-like main cable element (solid line), that divides the BADJ in two primary alveolar ducts, is clearly visible. In addition, the complexly structured helical line elements (dashed lines) are visualized, illustrating the true extent and 3D configuration of these complex structures.

**Figure 3.**
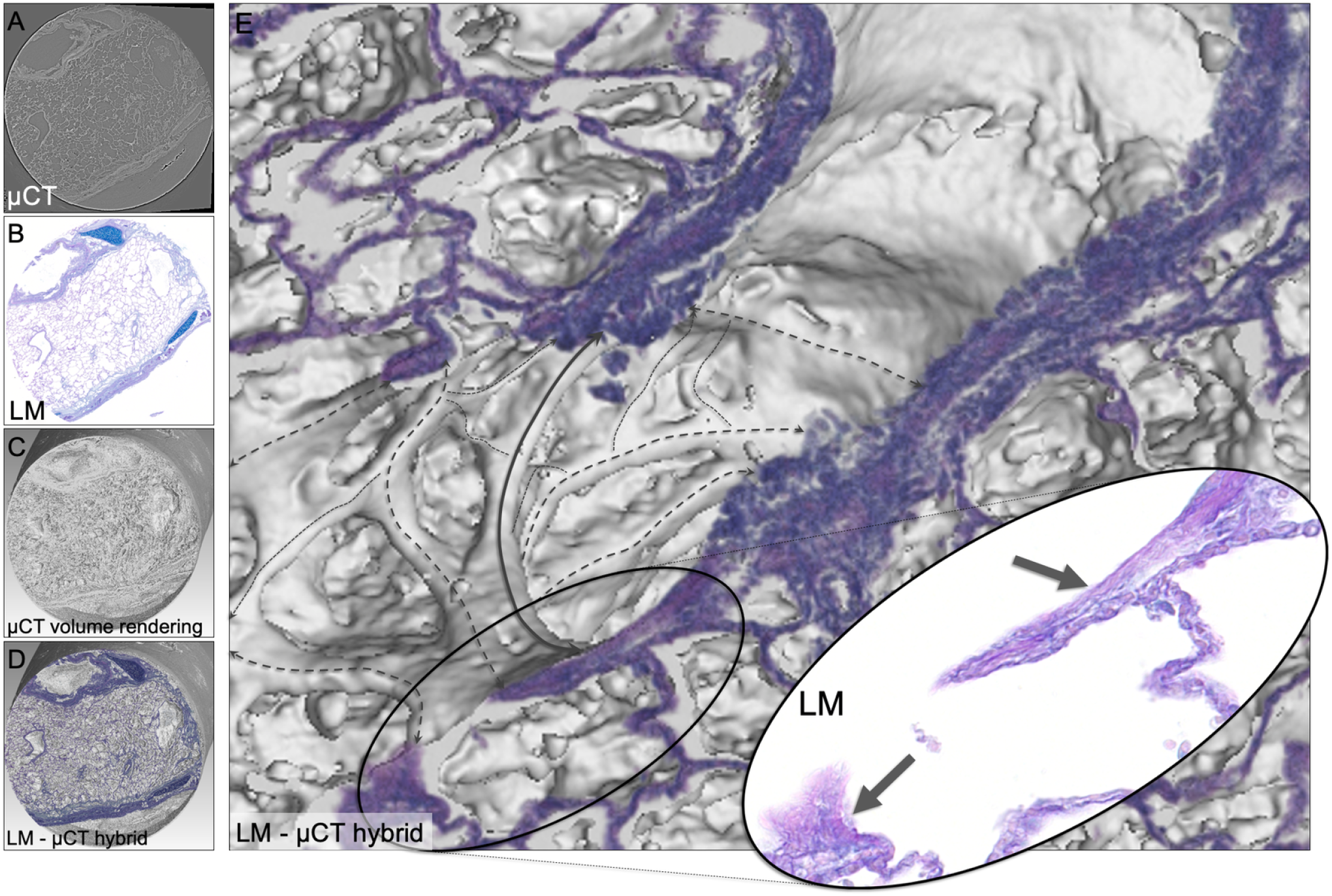
LM and SRμCT hybrid depiction of BADJ. A) exemplary slice from the reconstructed synchrotron radiation μCT data set of the tissue cylinder acquired via punch biopsy, B) digital light microscopy image (LM), AB-PAS stain. C) Volume rendering shows the cylindrical shape of the SRμCT data set. D) Hybrid image of the registered LM and the SRμCT data set. E) A magnified portion of the hybrid image shows a BADJ. The black arrows in the magnified LM insert indicate the alveolar septal tips, while in the CT data the complex 3D architecture of the axial connective tissue system is portrayed. A carina-like main cable element (solid line) divides the BADJ in two primary alveolar ducts. The complexly structured helical line element gives rise to alveolar entrance rings (dashed lines).

In classical LM the axial connective tissue system is inadequately represented due to its complex 3D structure (Fig. 4A). Only its 2D cross-sections are typically referred to as so-called septal tips (Fig. 3E detail view). The 3D scans provide sufficient contrast to allow for segmentation of the cable like structures that make up the axial connective tissue system, however a reliable and verified method is needed to securely classify their histological origin. As demonstrated in Fig. 4B, when an AB-PAS stained histological LM image is spatially registered to the SRμCT data sets, its specificity for connective tissue fibers can be used to generate an informed 3D segmentation of the axial connective tissue system from the integrated SRμCT data set (Fig. 4B yellow). Thereby, this essential anatomical structure proposed by Weibel et al. (Weibel, 2009) (Fig. 4A) can now be verified and visualized and pathological changes affecting its 3D integrity may be assessed. The information was further supplemented by spatially registered SEM which allowed to analyze the ultra-structure of the same septal tips found in the LM images (Fig. 4B detail view), via a down-stream component of the imaging pipeline outlined below. The specimen shown in Fig. 4B is a FFPE lung specimen from a human patient that died due to COVID-19.

**Figure 4.**
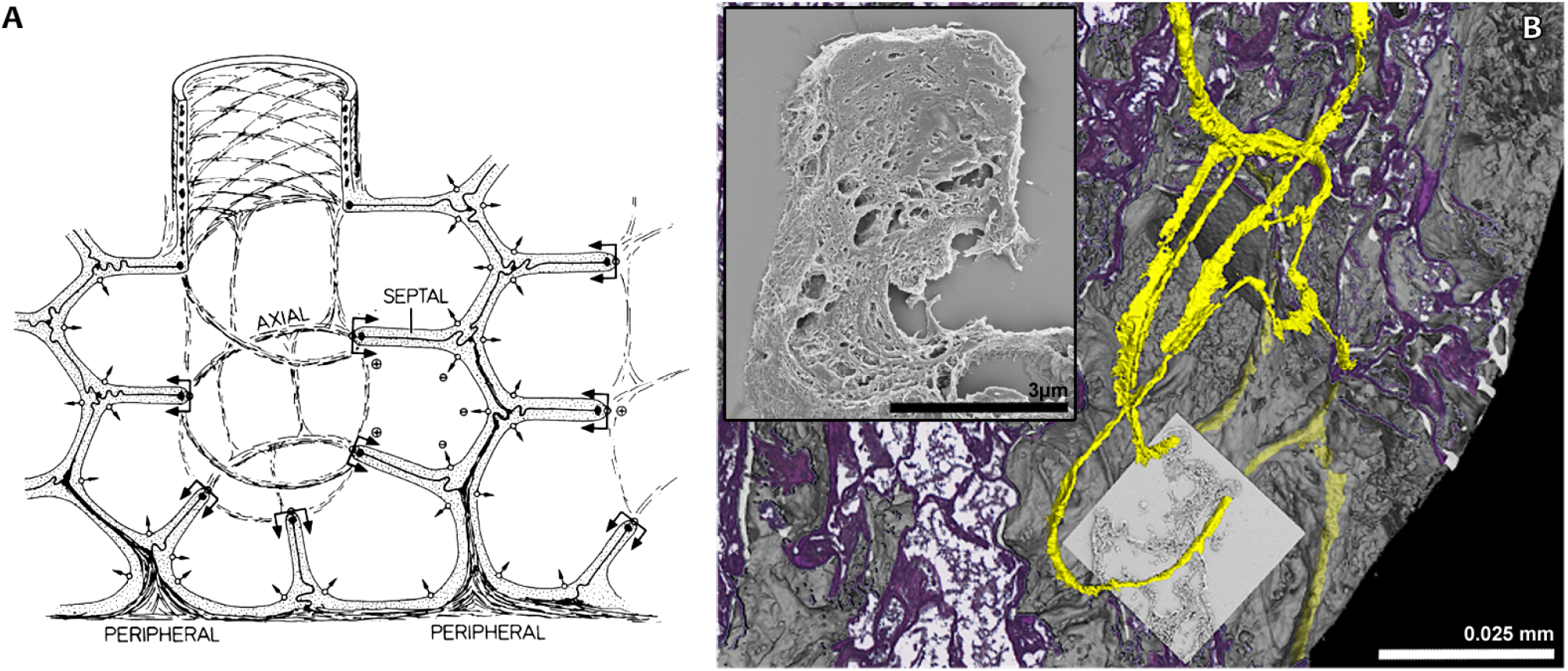
Multi-modal imaging validates 3D-structure of the alveolar duct fiber system. A) Schematic conceptual illustration of the axial connective tissue system of the lung by Weibel Weibel (2009). Alveolar entrance rings are kept under tension and held open by helical elastin cable elements. B) The combination of imaging modalities allows a detailed view of parts of the lung’s axial connective tissue system. The 3D rendering of a SRμCT scan of a FFPE lung specimen of a human patient that died due to COVID-19 has been virtually cut at the exact position of the histological section. Due to a detected difference in contrast, helical parts of the axial connective tissue system were successfully segmented with a region growing based tool. After segmentation, the helical structures can be traced inside the sample and can be viewed in their three-dimensional context, thus allocating them at the very edge of alveolar entrance rings. As confirmed by histological staining, these helical parts consist of elastin cable elements. Through targeted SEM, the ultrastructural surface composition of the elastin cable elements is shown.

### Same-slide scanning electron microscopy facilitates sub-micron correlative imaging montage

SEM supplements tissue analysis with information about the tissue’s ultra-structure and surface topography. Due to its large depth of field, the method yields a three-dimensional appearance useful for understanding the fine surface structure of the sample. Similar to μCT, the measurements cannot directly be assigned to different tissue types. Thus, the same strategy of combining it with histology as a tissue specific method is of great interest. For the mapping of structures depicted by SEM directly to their exact counterpart in light micrographs of histological samples, we applied same-slide SEM. Histological processing, staining, imaging and subsequent preparation for SEM as well as imaging was performed on conventional glass slides. This provided the best results possible for both imaging modalities. Between histological processing and sample preparation for SEM, no significant mechanical deformation occurred, resulting in images that were immediately fit for integration without prior elastic registration. Fig. 5A shows a H&Estained slice of the punched FFPE porcine lung tissue sample. Black rectangles indicate a cartilage region (Fig. 5B) as well as a lung parenchyma region containing septal tips (Fig. 5E). Fig. 5C and D show the respective SEM imaging results for ‘same-slide’ EM. The checkerboard visualizations in Fig. 5D and G show that LM and SEM data could be overlaid perfectly.

**Figure 5.**
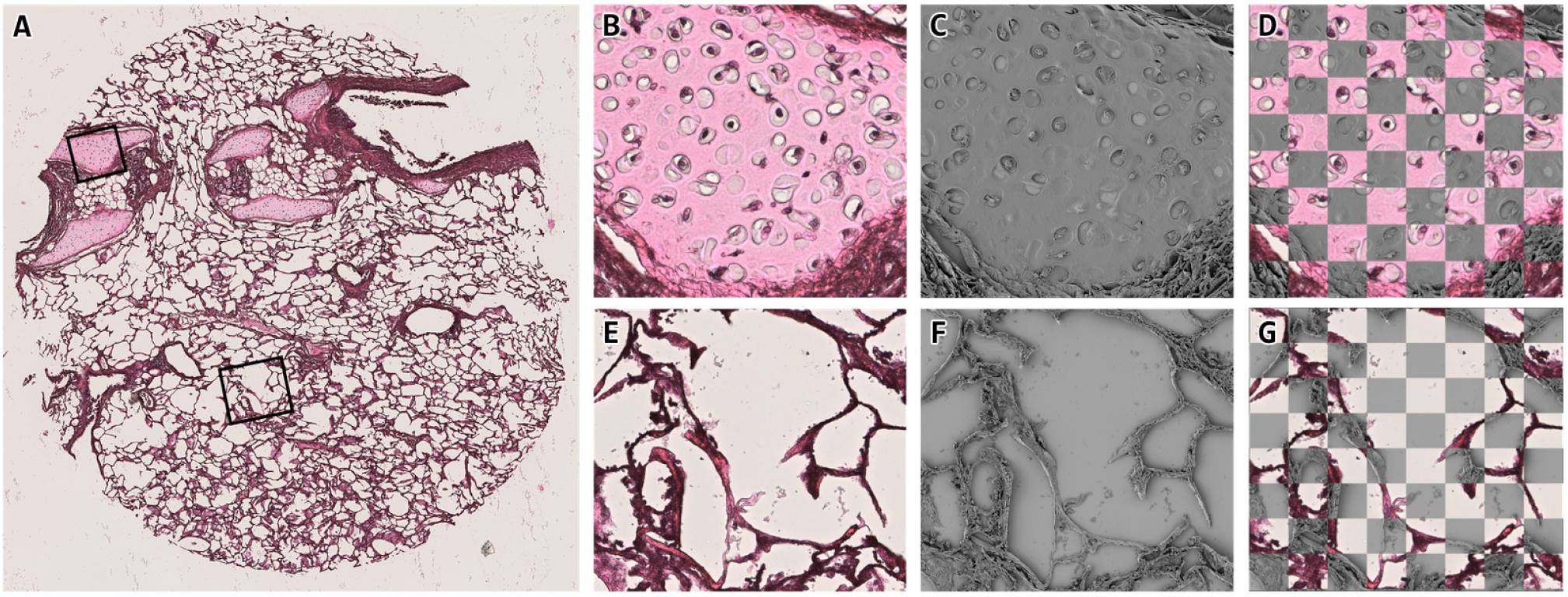
Elastic registration of histology and SEM. A) H&E-stained section of the analyzed 4 mm diameter tissue cylinder from the paraffin embedded lung tissue. Rectangles indicate two exemplary regions that have been analyzed by SEM. B) and C) show the same region imaged via histology and SEM respectively. D) a checkerboard visualization demonstrates that both data sets overlap perfectly. E) to G) show the same perfect overlay for a second region of interest. To achieve this overlap, no deformation needed to be compensated to match the data, demonstrating the most significant benefit of the same-slide SEM technique.

### Elastic image registration allows for creation of congruent multimodal hybrid images

As depicted in Fig. 4 same-slide SEM registers perfectly to prior acquired light microscopy images of the same tissue. Using the large field-of-view (FOV) of classical histology light microscopy imaging, the overview data can be registered into the 3D context of the SRμCT data set of the intact specimen. Elastic registration (Albers et al., 2021) was applied to compensate for mechanical deformations of the histological section that originated from the microtome sectioning process. Fig. 6A shows the 3D rendered SRμCT data set virtually cut open at the level of the corresponding histological section. The elastic registered histological slide as well as the SEM data were rendered into the same 3D context. To demonstrate the benefit of this approach supplementing histology with information on ultrastructural configuration via SEM and 3D information of the SRμCT acquisition the segmentation of a bronchus (blue), an arterial pulmonary blood vessel (red) and bronchial cartilage (pink) were displayed in 3D. The close-up in Fig. 6B demonstrates the precision of the registration and shows that now, the 3D information of the aforementioned structures can be related to their cross-sectional equivalent in histology as well as SEM.

**Figure 6.**
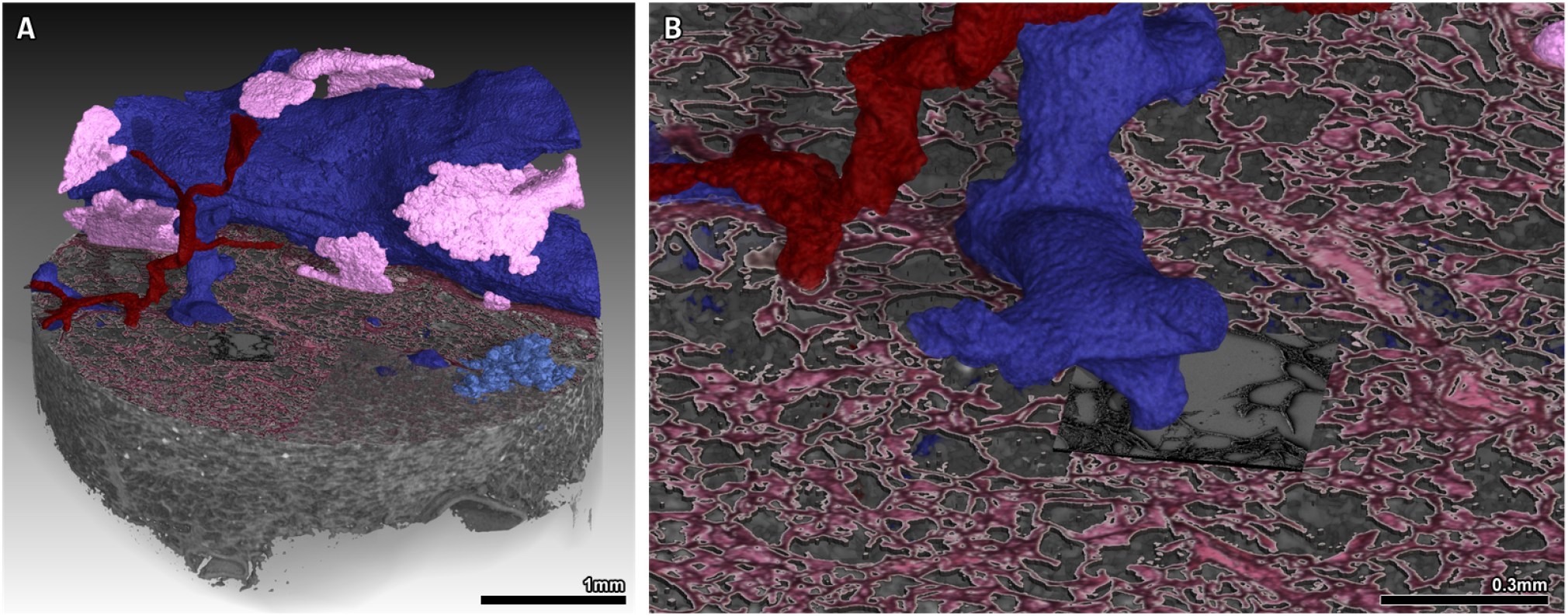
Hybrid image, created after elastic registration and subsequent integration of SRμCT, histology and SEM. A) Overview of all imaging modalities (SRμCT, H&E stained light microscopy and same-slide SEM) of the workflow combined into one. A bronchiolus with surrounding cartilage, a venous blood vessel and a terminal bronchiolus with its bronchoalveolar ductus and associated alveoli have been segmented. B) Zoomed in view of the hybrid image. Regions of interest for SEM, in this example containing septal tips, had been selected within the light microscopy image.

## Discussion

Classical histopathology, using light microscopy (LM) on formalin-fixed and paraffin embedded (FFPE) tissues, remains the gold standard for microstructural tissue analysis in biomedical research. However, its limitations in providing three-dimensional (3D) context and highresolution ultrastructural details present significant restrictions, particularly for complex organs like the lung. This study introduces an innovative integrative processing and imaging pipeline that combines synchrotron radiation based micro-CT (SRμCT), classical histology, and same-slide scanning electron microscopy (SEM) to overcome these limitations.

The incorporation of SRμCT into the pipeline provides a relevant advantage by offering volumetric imaging of entire paraffin blocks. This allows for a comprehensive 3D visualization of the whole tissue specimen and tissue architecture before any sectioning. Such prescreening capability is crucial for identifying regions of interest (ROIs) within the heterogeneous lung tissue, where pathological changes may be scattered and minute. The high-resolution SRμCT scans facilitate precise targeting of these ROIs for subsequent histological analysis, thereby enhancing the diagnostic accuracy and efficiency.

The study demonstrates that μCT images can effectively guide the microtome sectioning process, ensuring that specific planes of interest are accurately targeted. This is particularly beneficial for lung tissue, where maintaining the integrity of delicate structures like alveolar entrance rings and the axial connective tissue system is critical. By correlating SRμCT data with histological sections, the integrative pipeline maintains the spatial context of the tissue, providing a robust framework for detailed microstructural and possible further down-stream analysis. Histological analysis, with its various staining techniques, remains indispensable for identifying specific cell types and tissue components. However, its inability to provide ultrastructural details necessitate complementary techniques. The integration of same-slide SEM addresses this gap by offering ultra-high-resolution imaging of the targeted tissue’s ultrastructure. SEM’s capability to reveal fine surface details and sub-micron structures adds a critical layer of information, enhancing our understanding of fine structural components and the tissue’s microenvironment.

The study’s workflow ensures that μCT data, the histological and the SEM images are perfectly registered, allowing for seamless integration of data from all modalities. A key challenge in multimodal imaging is aligning data from different modalities due to variations in sample preparation and imaging processes. The study employs elastic image registration techniques to align histological slides with SRμCT and SEM data accurately. This approach compensates for deformations introduced during the sectioning process, ensuring that the spatial relationship between different datasets is maintained. Elastic registration not only enhances the accuracy of data integration but also allows for the creation of congruent hybrid datasets. These datasets provide a comprehensive view of the tissue, combining 3D structural information from SRμCT, specific histological details, and ultrastructural insights from SEM. This multimodal approach enables researchers to correlate histological features with their ultrastructural counterparts in a comprehensive 3D context, providing a holistic view of the tissue. For instance, the study highlights how SRμCT provides the 3D configuration of the axial connective tissue system and SEM can be used to examine the ultrastructure of septal tips in lung tissue, a complex structure that is challenging to resolve with LM alone. The integration is exemplified in the study by the detailed visualization of the broncho-alveolar-duct-junction (BADJ) and the axial connective tissue system, critical structures in lung tissue that are often inadequately represented in conventional histology.

For a comprehensive analysis, typically several methods are combined such as histology, genomics, proteomics and more. However, due to the fact that each of these modalities requires different tissue processing strategies, they are usually done on different parts of the specimen, which in turn limits the diagnostic value especially in diseases characterized by small lesions and a large tissue heterogeneity (Bingham et al., 2020). To this end spatially resolved omics techniques have been developed, which however lack the ability to observe a larger portion of the tissue in 3D.

Since any standard FFPE specimen can be used for label-free virtual histology by the means of phase contrast μCT as shown by multiple studies (D’Amico et al., 2024; Frost et al., 2023; Svetlove et al., 2023), the proposed method can be integrated into the standard workflow of classical pathological tissue analysis which is based on unstained FFPE tissue samples. This distinguishes the presented workflow from others that require more specific sample preparation in terms of e.g. staining and embedding (Bosch et al., 2022), thus not making them applicable to the aforementioned unstained FFPE tissue samples. As already demonstrated by for instance D’Amico et al. (D’Amico et al., 2024), our workflow can be applied to archived samples and extended by various methods for analysis of FFPE soft tissue like phase contrast CT, histology and atomic force microscopy. This reveals the huge potential scientific treasure on which research institutions and pathological institutes worldwide are sitting in the form of archived FFPE samples in their tissue banks, which have not yet been analyzed in three dimensions.

The synchrotron-based phase contrast SRμCT imaging method employed in this study is, in general, freely accessible to researchers after an experimental proposal have successfully been reviewed, for instance via the EuroBioImaging initiative www.eurobioimaging.eu. Nevertheless, the access to synchrotron light sources will continue to be limited. Here we showed that also an X-ray microscope (Zeiss 620 Versa) can be utilized in a laboratory setting and potentially allows to exploit phase contrast as for instance demonstrated by Bidola et al. (Bidola et al., 2015). In our case, the data sets obtained with the data produced by Versa 620 showed even better image quality due to the higher spatial resolution. However, even considering more efficient setups or parameter settings for laboratory based X-ray microscopy, the shorter scanning time to acquire SRμCT data is making very rapid acquisitions of larger cohorts of specimens possible as for instance presented by Albers et al. Albers et al. (2024). Nevertheless, the ability to perform our workflow independently from a synchrotron facility underlines its general and wide applicability.

In both cases, high resolution μCT imaging will generate a huge dose within the specimen. So far, we did not observe any deterioration of the FFPE tissue or a reduction in image quality during the histological analysis. However, we observed an increase in brittleness, which rendered sectioning of the FFPE blocks more challenging after SRμCT scans. The integration of additional modalities for downstream analysis of tissue sections into the 3D context is a promising area for further exploration. Advanced imaging techniques, such as multiplexed fluorescence imaging and superresolution microscopy, could provide more detailed insights into molecular and cellular interactions within tissues. Additionally, computational methods like machine learning and artificial intelligence could enhance image analysis, enabling automated identification and quantification of pathological features across different scales.

## Conclusion

The execution of the workflow is demonstrated on unstained and otherwise unmodified FFPE tissue samples, the standard method for embedding and archiving samples in anatomy and pathology. This implies that the workflow can be applied to any FFPE specimen that has already been processed and archived in the past. The presented pipeline has a huge potential for machine learning based projects in digital pathology such as shown by Song et al. (Song et al., 2023) which profit from a database as big and diverse as possible. Advancements in integrative tissue analysis methodologies are crucial for overcoming the limitations of traditional histological techniques. By combining multiple imaging modalities and preserving the 3D context of tissue structures, researchers can gain a more comprehensive understanding of tissue architecture and pathology. This is particularly important for organs like the lung, where complex structural integrity is vital for function and disease progression often involves heterogeneous and scattered alterations. The development and refinement of such integrative approaches may continue to enhance our ability to diagnose and treat diseases with greater precision and efficacy.

## Methods

The imaging pipeline described in the following paragraphs was developed using a porcine lung due to its similarity in size and macroanatomy to a human lung (Judge et al., 2014). FFPE lung specimens were μCT-scanned using a synchrotron radiation-based and a commercially available laboratory system. Targeted punch biopsies were taken from regions of interest and imaged at higher resolution. After examination of the biopsy cylinder scans for structures of interest, pre-selected tissue slices (5 μm thickness) were taken from targeted planes of interest, placed on conventional glass slides, deparaffinized, stained and imaged using digital LM. The same slice was further processed and SEM was performed on regions pre-defined in the LM scan. Fuxlastix was applied to register the LM and SEM tissue scan to the SRμCT data to correct for deformation from LM/SEM preparation. The multi-scale and multi-modal image data was integrated, depicted and analyzed in VGStudio Max. The workflow is shown in Fig. 7.

**Figure 7.**
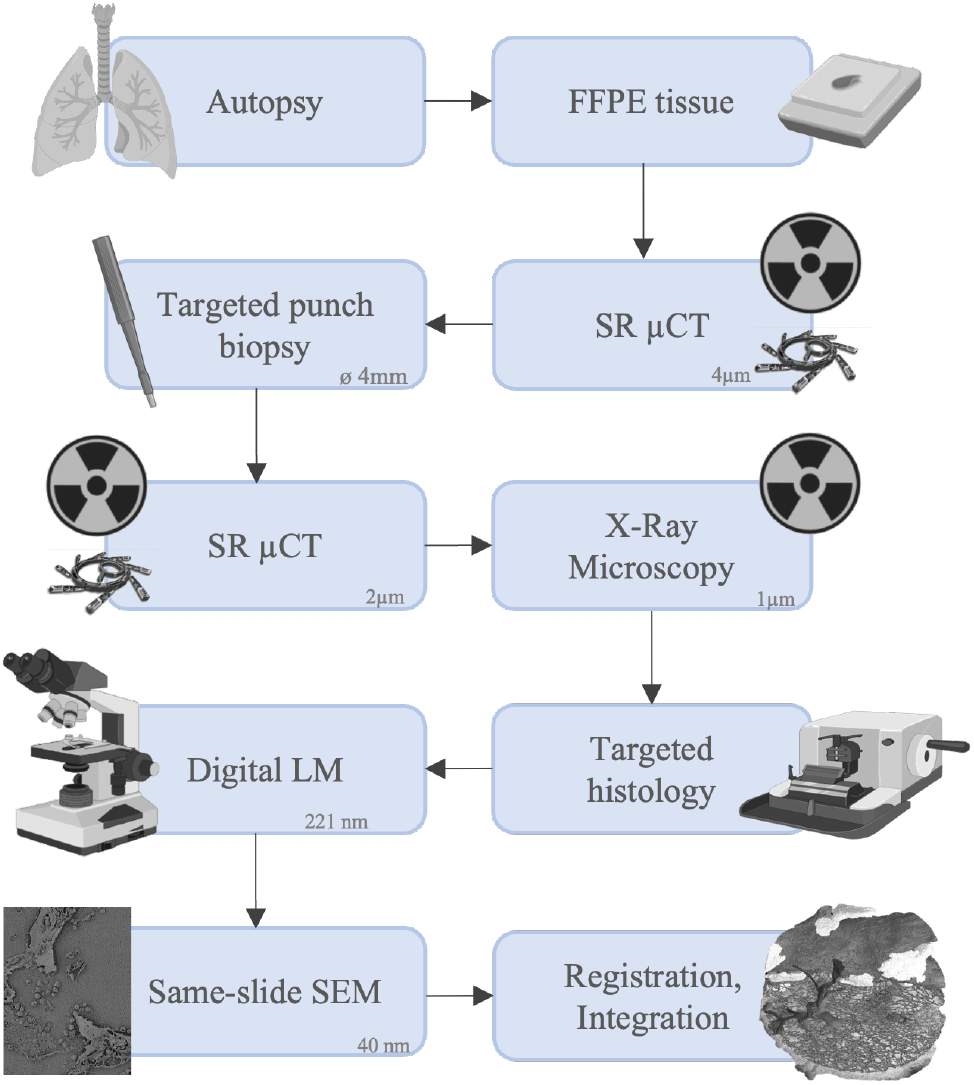
Schematic depiction of the workflow that has been developed and applied successfully. During macroscopic sampling, lung tissue blocks were collected and afterwards formalin fixed and paraffin embedded. SRμCT scans were obtained at a voxel size of 4 μm. The scans were examined, regions of interest were selected and a targeted punch biopsy was taken, yielding a tissue cylinder with a diameter of 4 mm. The tissue cylinder was again scanned at a voxel size of 2 μm. The same tissue cylinder was also scanned in a conventional X-ray microscopy setup at a voxel size of 1 μm. The cylinder scans were evaluated for structures of interest. Targeted histology was performed, histological sections were stained and digital LM images with a pixel size of 221 nm were produced. On the same glass slide, the stained histological sections were prepared for SEM, and SEM images were taken, concluding the imaging pipeline with the highest acquired pixel size of 40 nm. Ultimately, all positions where imaging modalities had been performed were relocated within the 3D rendering of the scanned volume. Elastic registration was performed to enable integration of all imaging modalities.

### Synchrotron phase contrast CT (SRCT)

The specimens were scanned at the SYRMEP beamline (SYnchrotron Radiation for MEdical Physics) at the Italian synchrotron “Elettra” (Trieste, Italy) using the white beam high resolution setup for specimens as described by Dullin et al. (Dullin et al., 2021). We used a pixel size of 4.5 μm and a sample-to-detector distance of 45 cm. 1800 projections with an exposure time of 50 ms were acquired over 180°. Given the relatively small lateral beam size of approx. 3.5 mm up to 3 overlapping scans with an increment of 3 mm were performed. In addition, a 4 mm diameter tissue cylinder was obtained via punch biopsy and scanned with a sample-to-detector-distance of 150 mm, a voxel size of 2 μm and an exposure time of 100 ms. Single distance phase retrieval (homogeneous version of transport of intensity equation, TIE_HOM (Paganin et al., 2002)) with a delta-to-beta ratio of 200 was applied prior to 3D reconstruction using the classical filtered back projection both implemented in the STP-Gui project (Brun et al., 2015).

### Zeiss X-ray microscopy

For further image data acquisition a Zeiss 620 Versa was used. A region of interest was identified within the tissue volume and detail scan with 1 μm voxel size, 3 s exposure time, 4x objective lens, 40 kV tube voltage and 5001 projections over 360°was performed, resulting in a total scanning time of 9.3 hrs.

### Re-embedding and histological sectioning

The 4 mm tissue cylinder was re-embedded in paraffin to enable microtome-based histological sectioning. After calculating the cutting depth required to hit the predefined targeted zone, sections were taken at a thickness of 5 μm. Starting from the beginning of the targeted zone, sections were collected, placed on conventional uncoated glass slides and processed further.

### Staining / Light microscopy

After targeted specimen collection, the tissue sections were deparaffinized and rehydrated. For this sample, H&E and AB-PAS staining were applied and cover slips were added. The light microscopy images were then acquired at a resolution of 221 nm/px using Hamamatsu C13210. Examination of the images was carried out with the software NDP.view2 and QuPath V.0.3.2.

### Scanning electron microscopy [SEM]

The cover slips of the histological section were removed in xylol. The specimen were then sputtered with 3 nm Pt/Pd in a relation of 80/20 with Quorum A150V ES plus (clean current 50 mA, sputter current 5 mA). The slides were placed on a carousel and connected with copper tape (Plano GmbH, G3490) and silver conductive fluid (Plano GmbH, ACHESON 1415, G3692) to prevent charging effects. Navigation to specific sites of interest was performed using the Atlas 5 software (ZEISS). SEM acquistion was performed with a Ultra TM 55 (Zeiss) using 2 kV voltage and a 40 nm pixel size SE detector.

### Elastic image registration and quality assessment

2D-3D elastic registration was achieved using Fuxlastix (Albers et al., 2021), a front-end to Elastix (Klein et al., 2009; Shamonin et al., 2014). Given the different image content of CT, histology and SEM normalized mutual information was used as cost function. Due to the non-uniform deformations in the cutting process a grid of cubic thin-plate b-splines was used as deformation field.

### Statistical methods and used software

VGStudioMAX (VolumeGraphics, Heidelberg) was used for rendering and 3D tissue analysis. Fuxlastix, a front-end to Elastix, was used to perform 2D-2D elastic registration as described in Albers et al. (Albers et al., 2021). Custom python scripts were used to calculate the above mentioned quantitative measures utilizing standard python modules such as numpy, scikit-image and matplotlib.

### Ethical statement

#### Human material

The study was approved in advance by the Ethics Committee of the Medical Faculty of the University of Heidelberg (No. S-242/2020). All described studies on humans or human tissue were conducted with the approval of the responsible ethics committee, in accordance with national law and in accordance with the Declaration of Helsinki of 1975 (in the current, revised version).

#### Animal material

No animal was sacrificed for the specific purpose of this study, as the thoracic visceral organ block is an animal by-product of commercial meat processing. The preparations were carried out under regular veterinary supervision of an approved meat processing plant and were treated with the same hygiene measures as for fresh meat. In accordance with Article 23 of Regulation (EC) 1069/2009, registration for the storage, handling, transportation and disposal of animal byproducts for research purposes in category 3 was registered to the competent veterinary regulatory authority under registration number DE08221103321.

## Acknowledgements

The authors would like to thank Jolanthe Schatterny (TLRC Heidelberg) for processing and providing the histological specimens included in this study.

I.W., R.C. and R.R.S. acknowledge funding from the German Federal Ministry of Education and Research, project NanoPatho, FKZ 13N14476 and by the German Research Foundation EXC 2082/1-390761711 (3D Matter Made to Order).

Figure 7 was created with Biorender.com.

